# Full structural ensembles of intrinsically disordered proteins from unbiased molecular dynamics simulations

**DOI:** 10.1101/2020.06.16.155374

**Authors:** Utsab R. Shrestha, Jeremy C. Smith, Loukas Petridis

**Author notes:** This manuscript has been authored by UT-Battelle, LLC, under contract DE-AC05-00OR22725 with the US DOE. The US government retains and the publisher, by accepting the article for publication, acknowledges that the US government retains a nonexclusive, paid-up, irrevocable, worldwide license to publish or reproduce the published form of this manuscript, or allow others to do so, for US government purposes. DOE will provide public access to these results of federally sponsored research in accordance with the DOE Public Access Plan (https://www.energy.gov/downloads/doe-publicaccess-plan).

## Abstract

Molecular dynamics (MD) simulation is widely used to complement ensemble-averaged experiments of intrinsically disordered proteins (IDPs). However, MD often suffers from limitations of inaccuracy in the force fields and inadequate sampling. Here, we show that enhancing the sampling using Hamiltonian replica-exchange MD led to unbiased ensembles of unprecedented accuracy, reproducing small-angle scattering and NMR chemical shift experiments, for three IDPs of variable sequence properties using two recently optimized force fields. Surprisingly, we reveal that despite differences in their sequence, the inter-chain statistics of all three IDPs are similar for short contour lengths (< 10 residues).

## INTRODUCTION

Intrinsically disordered proteins (IDPs) exhibit biological function without folding spontaneously into a unique three-dimensional (3D) structure.^1^ IDPs are abundantly present in all proteomes and play major roles in signaling, transcriptional regulation and regulation of phase transitions in the cell via processes that may involve high-specificity or low-affinity interactions and recognition of partners by folding upon binding.^1–5^ About 50 to 70% of the proteins in the human genome associated with cancers, diabetes, cardiovascular, and neurodegenerative diseases have a minimum of 30 residues that are intrinsically disordered, making IDPs possible drug targets.^1^ Additionally, IDPs are an essential part of plant immune signaling components and also mediate plant-microbe interactions.^6^

Understanding the function of a protein requires a determination of its 3D structure.^7^ IDPs adopt highly dynamic structural ensembles, which are commonly characterized by nuclear magnetic resonance (NMR)^8^, small-angle X-ray/neutron scattering (SAXS/SANS),^9,10^ singlemolecule Förster resonance energy transfer (smFRET),^11^ hydrogen-exchange mass spectrometry^12^ and circular dichroism (CD).^13,14^ However, the information content of the applied experimental techniques is insufficient to obtain the ensemble of 3D conformations an IDP adopts.^15^ The experimental observables often represent averages over the ensemble and the data are typically sparse, providing too little information to unambiguously determine the 3D ensemble.

Molecular dynamics (MD) simulation can in principle provide the missing information and furnish a complete atomic resolution description of IDP structure and dynamics.^2^ Recent optimizations of the protein and water potential energy functions^2,16–27^ have led to accurate simulation of short disordered peptides and model systems.^17,18,28–31^ However, the simulations are not always consistent with experiment either because of inadequate sampling or shortcomings of the force fields.^2,18,21,23,29,32,33^

A common and successful approach to determine an IDP configurational ensemble is to force the MD results to match existing experiments, either by biasing the MD potential,^34,35^ or by *a posteriori* reweighting the ensemble of the MD population.^36,37^ One challenge for these methods is degeneracy, *i.e*. distinct 3D conformations may yield the same observable, which may lead to over-fitting. Bayesian maximum entropy optimization approaches, which aim to perturb the MD ensemble as little as possible, have been employed to avoid over fitting.^34,37,38^ However, these approaches always require a prior experimental measurement and do not afford a predictive understanding of IDPs.

Recently, by enhancing the configurational sampling of MD simulations using Hamiltonian replica-exchange MD (HREMD) the configurational ensemble of an IDP was generated that is in quantitative agreement to SAXS, SANS and NMR measurements without biasing or reweighting the simulations.^39,40^ HREMD improves sampling by scaling the intraprotein and protein-water potentials^16,19^ of higher-order replicas, while keeping the potential of the lowest rank replica unscaled^41–44^ so as to represent the physically-meaningful interactions of the system. However, two IDPs^39,40^ were studied and the general applicability of this approach has not been established.

Here, we report that HREMD produces configurational ensembles consistent with SAXS, SANS and NMR experiments for three IDPs with markedly different sequence characteristics: Histatin 5 (24 residues) and Sic 1 (92 residues), both of which have an abundance of positively charged residues, and the N-terminal SH4UD (95 residues) of c-Src kinase, which contains positively and negatively charged residues mixed over the sequence. The HREMD results are in agreement with experimental data on both local and global properties when employing either of two force fields (Amber ff03ws^19^ with TIP4P/2005s^19^ and Amber ff99SB-*disp*^16^ with modified TIP4P-D,^16^ hereafter termed as a03ws and a99SB-disp respectively). In contrast, standard MD simulations of equivalent computational cost as HREMD generate ensembles consistent only with NMR, but not with SAXS. Further, the HREMD ensembles of IDPs are found to be described by a theoretical semiflexible polymer chain model quantifying the stiffness and strength of interaction with solvent. We suggest “best practices” in achieving accurate and efficient IDP sampling using HREMD and discuss differences in the size between Sic 1 and SH4UD. The results demonstrate quite clearly that the recently optimized force fields are reliable and that the current major impediment to accurate simulation of IDPs using is sampling. HREMD is therefore the present tool of choice for obtaining atomic-detailed IDP ensembles.

## RESULTS

### HREMD ensembles in agreement with SAXS, SANS and NMR

We conducted HREMD simulations of three IDPs with varying amino acid composition (**Fig. S1**), employing two force fields: a03ws^19^ and a99SB-disp^16^. For comparison, we also conducted standard MD, *i.e*. without enhancing the sampling, of the same cumulative length as the HREMD (**Tables S1-S4**). The histograms of a radius of gyration (*R_g_*) show the IDPs adopt a continuum of collapsed to extended structures (**Fig. 1a-c**).

**Fig. 1.**
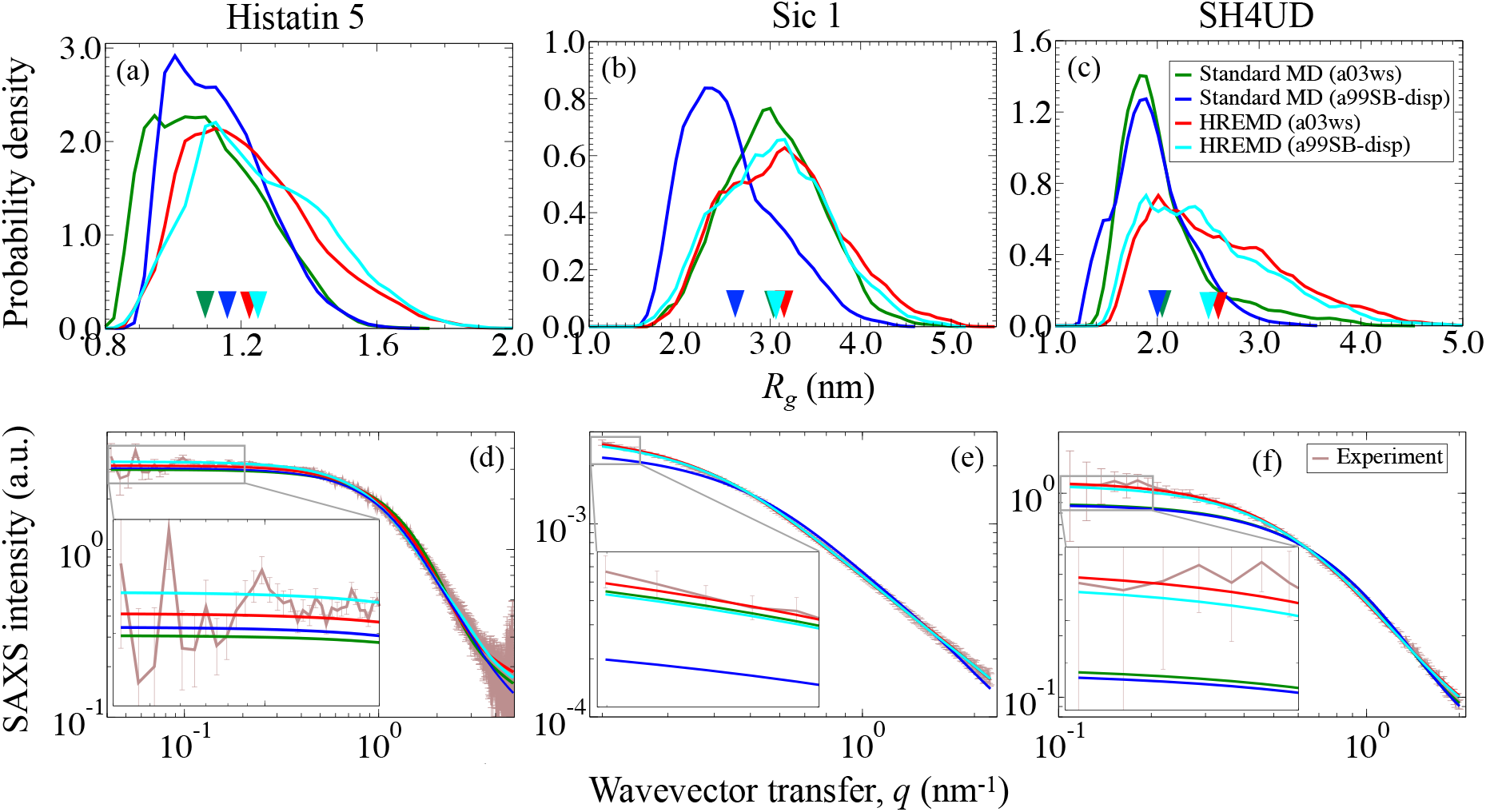
(a-c) The histograms of *R_g_* of (a) Histatin 5, (b) Sic 1 and (c) SH4UD obtained from MD simulations. The inverted triangles indicate the average *R_g_* of each simulation. (d-f) The SAXS profiles calculated from simulations (using SWAXS^45^) are compared to experiments for (d) Histatin 5,^30^ (e) Sic 1,^46^ and (f) SH4UD.^40^ Insets: SAXS data are zoomed at low-*q* values to show the differences in intensity for different force fields and sampling methods. In all cases the color code indicates the force fields, a03ws^19^ or a99SB-disp,^16^ and sampling methods, standard MD or HREMD (**Tables S1 and S2**). HREMD results are from the lowest rank replica of the simulations shown by bolditalics font in **Table S2**. SANS data of SH4UD are shown in Fig. S2.

The global, ensemble-averaged properties of IDPs such as *R_g_*, shape, and chain statistics can be derived using small-angle scattering. We calculated the ensemble-averaged theoretical SAXS and SANS curves from the simulation trajectories, by taking into account explicitly the protein hydration shell and without reweighting, and compared them directly to the experiments. We found an excellent agreement of the HREMD-generated ensembles with SAXS and SANS measurements for both force fields (SAXS in **Fig. 1d-f** and SANS in **Fig. S2**), whereas the standard MD simulations were found to deviate from the experiments, except for Sic 1 with a03ws. The agreement between simulation and experiment was quantified with the *χ*^2^ value as defined in **Eq. (5)** and listed in **Table S5**. The histograms of *R_g_* show that standard MD simulations sample more compact structures than does HREMD with the same force fields. Therefore, for the IDPs studied here, poor agreement with experiment arises primarily from insufficient sampling rather than from shortcomings of the force fields.

NMR chemical shifts provide information on the local chemical environment of protein atoms and reflect structural factors such as backbone and side-chain conformations. To further validate the simulations, we calculated the ensemble-averaged backbone chemical shifts (*C^α^, C^β^* and *N^H^* for Sic 1 and SH4UD, and *H^N^* and *H^α^* for Histatin 5) and compared to previously reported experiments (**Figs. 2, S3-S6**). The agreement between theoretical and experimental NMR chemical shifts was quantified by calculating the mean normalized deviation as defined by **Eq. (6)**. For Sic 1 and SH4UD, we found an excellent agreement with the experiments for all force fields and sampling methods, whereas the agreement is not quite as good for Histatin 5 (**Figs. 2 and S3-S6**).

**Fig. 2.**
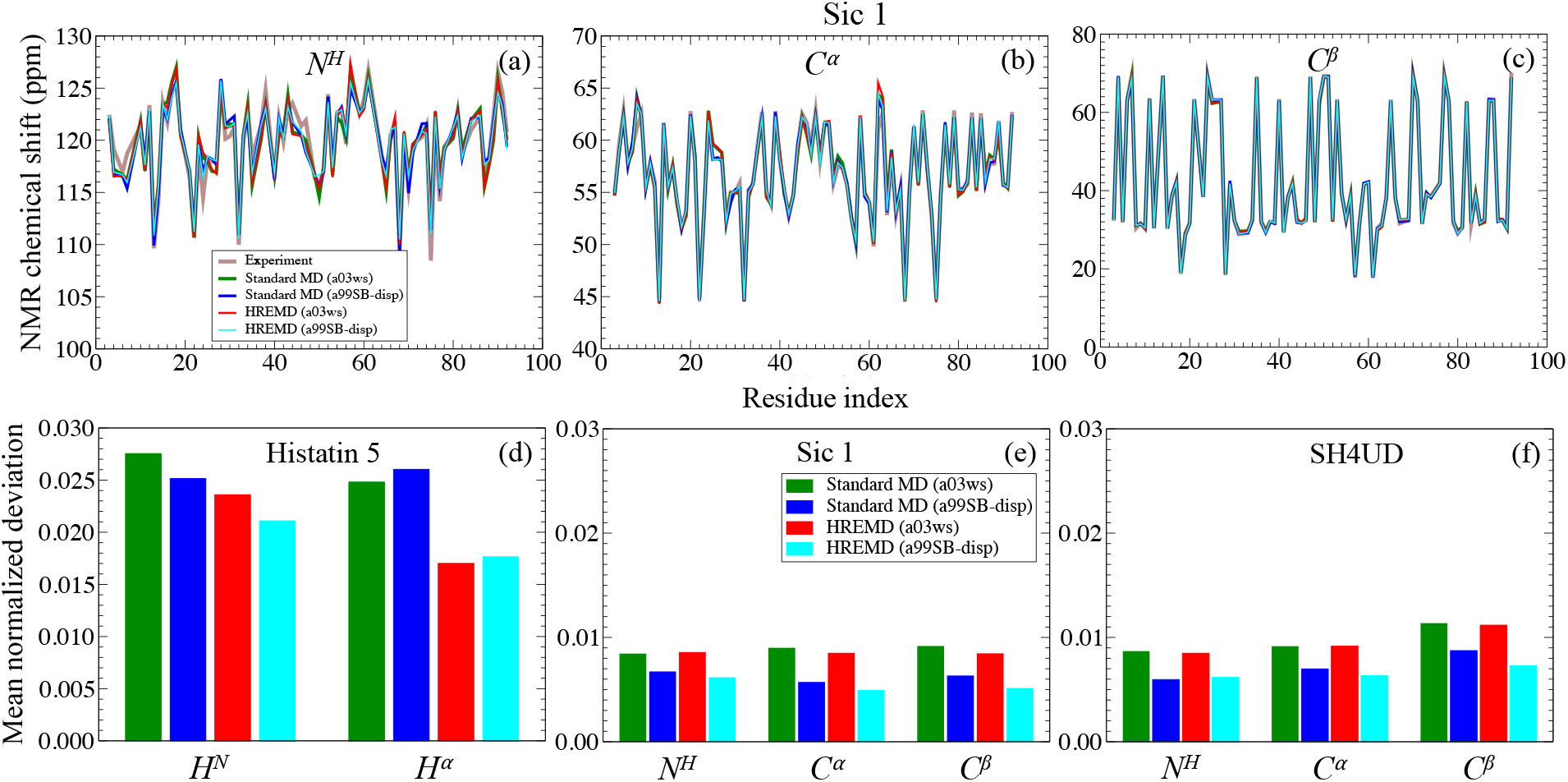
Comparison between the ensemble-averaged experimental and calculated NMR chemical shifts of backbone atoms (a) *N^H^*, (b) *C^α^* and (c) *C^β^*, for Sic 1. The mean normalized deviation of MD-derived NMR chemical shifts of backbone atoms with respect to experimental values, as defined in **Eq. (6)**, for (d) Histatin 5,^47^ (e) Sic 1,^46^ and (f) SH4UD.^48^ The color code indicates the force field and sampling method used. The theoretical NMR chemical shifts are calculated using SHIFTX2,^49^ which has relatively high values of root mean square errors of 0.1711 ppm and 0.1231 ppm for *H^N^* and *H^α^* respectively compared to *N^H^, C^α^* and *C^β^*.

Both force fields and sampling methods predict nearly the same transient secondary structure elements. Transient helices, which are considered to be biologically relevant,^50–52^ were found proximal to known phosphorylation residues of Sic 1^46^ and to known lipid-binding or phosphorylation residues in SH4UD.^48,53^ In contrast, the propensity of each secondary structure element is found to depend on both the force fields and sampling methods (**Figs. S7 and S8**). The IDPs we studied mostly showed a high propensity for coils that lack secondary structure, consistent with the lack of long-range contacts found in the simulations (**Fig. S9**).

### Polymer properties

We estimated the stiffness of the protein backbones by calculating the orientational correlation function

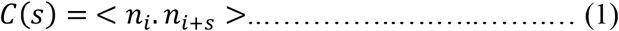

where *s*=|*i-j*| is the pairwise residue separation (sometimes called contour length), and *n_i_* is the unit vector connecting the backbone atoms N and C of residue *i*. The steeper the decay of *C(s)*, the lower the stiffness of the chain. *C(s)* is similar for the three IDPs for *s*≤10, exhibiting an exponential decay *C(s)* = *e^−s/l_p_^*, where *l_p_* is the persistence length. *l_p_* provides the maximum size of a protein segment over which the structural fluctuations are correlated. In other words, it is the measure of stiffness of a polypeptide chain. We found *l_p_* ~ 1 nm for all IDPs, in good agreement to the values for intrinsically disordered proteins.^54,55^ A power law decay (~*s^-3/2^*) is found for Sic 1 at 10<*s*≤26, whereas correlations decay more rapidly and vanish for *s*>10 for Histatin 5 and SH4UD. Therefore, Sic 1 is the stiffest.

The statistics of internal distances (“scaling properties”) of polymers in dilute solution can be characterized using the Flory scaling law given by Eq. (2):

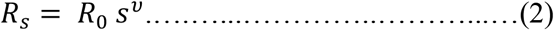

where *R_s_* is the average intraprotein pairwise distance between the C_α_ atoms of residues *i* and *j* at pairwise separation *s*=|*i-j*|, the prefactor *R*_0_ is a constant and *ν* is the Flory exponent. Balanced polymer-solvent and intrapolymer interactions give rise to Gaussian coil and *ν*=0.5, while a self-avoiding random walk (SARW) with *ν*==0.588 is predicted when the polymer-water interactions are favored. Interestingly, we found two different power law regimes are needed to fit the data^56,57^ (**Fig. 3b-d**). At short contour lengths (*s*≤10), *R_s_* is similar for all three IDPs, with *ν* ≈ 0.70, which indicates chain configurations stiffer than a SARW, and with a prefactor of *R_0_* ~0.4 nm (*R_0_* is the average distance between two consecutive C-alpha atoms). On the other hand, at longer residue separations (*s*>10) the *R_s_* of the three IDPs deviate. Histatin 5 and SH4UD with *ν* ≈ 0.43 and 0.40, respectively, adopt more collapsed global conformations than SARW. In contrast, Sic 1 (*ν* ≈ 0.60) remains stiff even at longer residue separations.

**Fig. 3.**
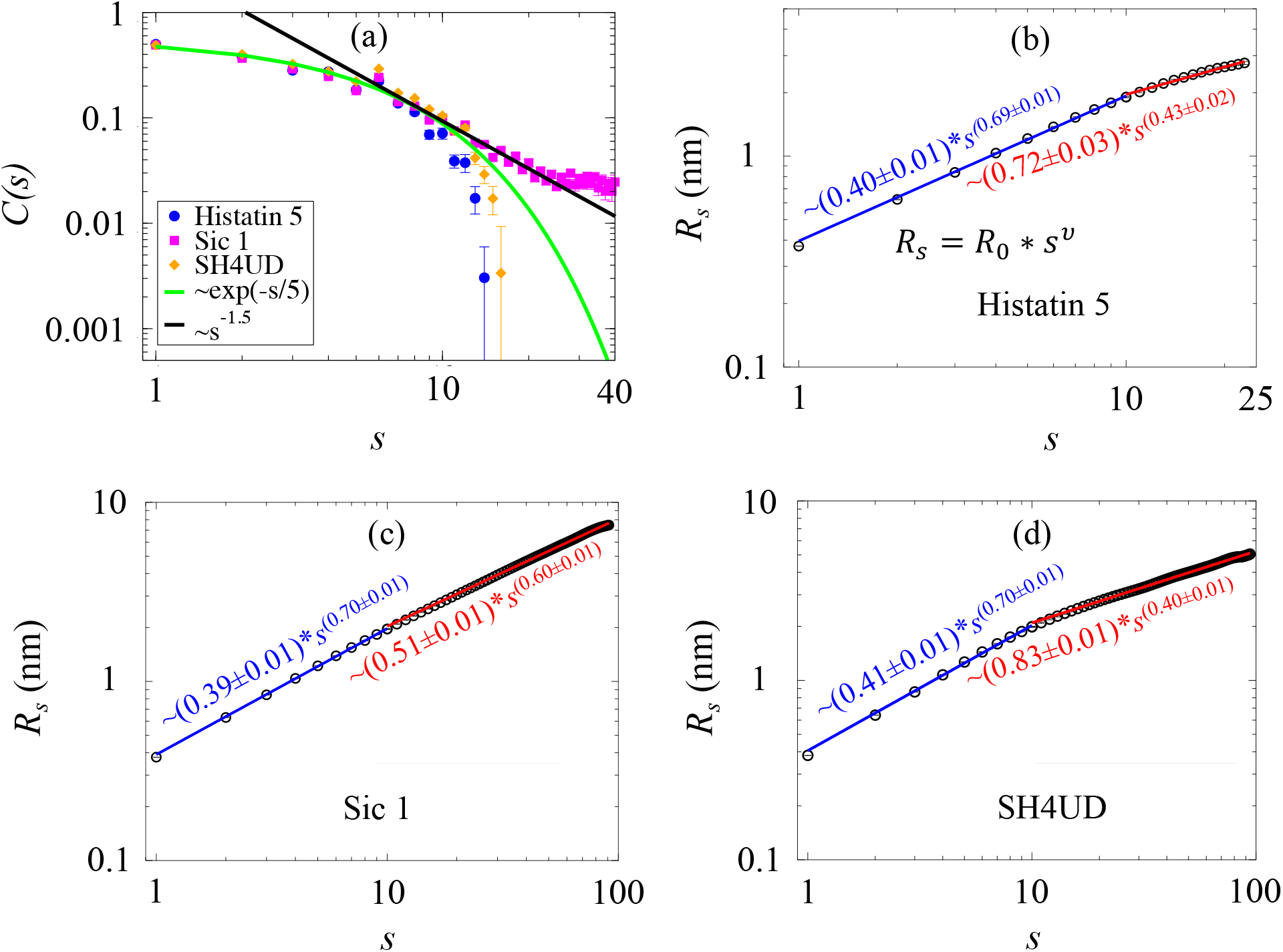
Chain statistics of IDPs. (a) The orientational correlation function as a function of the pairwise residue sequence separation, *s*. For *s* ≤10, *C(s)* is fitted by *C(s)* = *e^−s/l_p_^* for each IDP, which estimates the persistence length (*l_p_*). For *s*>10 the power law *C(s)~s*^-3/2^ applies only for Sic 1, whereas for Histatin 5 and SH4UD the correlation vanishes. (b-d) The average pairwise geometric distance (*R_s_*) between C-alpha atoms of two residues at separation *s* for (b) Histatin 5, (c) Sic 1 and (d) SH4UD. The data are fitted by Eq. (2) in two regimes, *s*≤10 (blue) and *s*>10 (red). The error bars are smaller than the symbol size.

## DISCUSSION

IDPs present a new paradigm for understanding flexibility-function relationships in biology.^1,58–60^ Currently, it is not possible to determine the ensemble of the 3D structures that an IDP adopts from either experiment or simulation alone. The number of experimental observables is considerably smaller than the number of the IDP’s configurational degrees of freedom, making model reconstruction from experimental data a highly underdetermined problem. For MD simulations, although improved molecular mechanics methods perform well for small model disordered peptides^2,18,28,30,31^, it has often been necessary to bias or reweight the MD results to achieve consistency with experiments.^34,35,37,38,61–63^ The reason MD has not always been accurate is unclear: it could be deficiencies in the force fields, insufficient sampling, or both.

Here, we demonstrated that HREMD reproduces key experimental observables (SAXS, SANS and NMR) using two different force fields for three different IDPs. In contrast, the ensemble generated by standard MD of equivalent length failed to match SAXS data (**Fig. 1**). The comparison of standard MD and HREMD using the same force field suggests the a03ws and a99SB-disp force fields are of adequate accuracy and that enhanced sampling techniques are necessary to reproduce experimental data.

We found that the calculated NMR chemical shifts and the *loci* of secondary structure elements are the easiest to converge as they are consistent between all the simulations, independent of force field and sampling method. In contrast, HREMD is required for SAXS observables to converge to the experimental values. The most difficult quantities to converge are the secondary structure propensities, which were found here to depend on both the force field and the sampling method, perhaps more on the former than the latter (**Fig. S7 and S8),** with a03ws and a99SB-disp having biases towards helices and β-sheets, respectively.

The data show that MD simulations can be in *apparent* agreement with NMR chemical shifts, which measure local structural information,^64^ while failing to reproduce SAXS/SANS intensities, which determine with high precision more global structural properties (here distributions of distances between pairs of nuclei that are more than ~1 nm apart^61,65^) (**Figs. 1, 2 and S10**).^40^ Thus, agreement with NMR alone is not always a definitive test of the accuracy of MD simulations of IDPs. It is critical to analyze and compare both local and global properties^16,66^ of IDPs to ensure the simulations have indeed generated accurate ensembles.

Simple theories established for semiflexible homopolymers and heteropolymers have been shown to provide a qualitative description of IDP structural properties such as stiffness^67–69^ and solvent quality.^11,13,70–72^ The high fidelity HREMD trajectories reveal that, despite having markedly different sequences, the IDPs studied here have a common hierarchical chain architecture. For short contour lengths (up to ~10 residues) the chain statistics of all three IDPS are similar, as evidenced by *R_s_* and *C(s)*. These short segments are relatively stiff with a Flory exponent of ν~0.7. Beyond this critical contour length, the IDPs differentiate. SH4UD and Histatin 5 become flexible, while Sic 1 remains relatively stiff with power-law decay in *C(s)* that implies long-range spatial correlations.^68^ This is consistent with Sic 1 being more extended than SH4UD.

The origin of the stiffness of Sic 1 relative to SH4UD can be understood by examining their primary sequences (**Fig. S1**). All the charged residues of Sic 1 are positive, leading to electrostatic repulsion between them. Further, Sic 1 contains 15 proline and 5 glycine residues. Proline is stiff due to its cyclic sidechain, whereas the absence of a sidechain for glycine increases backbone flexibility, which is known to be disorder-promoting.^55,73^ In comparison, SH4UD has both positively and negatively charged residues, 11 prolines and 12 glycines.

We now discuss the HREMD method^41,42,44^ and make recommendations for its optimal use in IDPs. HREMD enhances sampling by changing the quality of water as a good solvent for an IDP. This is achieved by effectively heating up only the solute by scaling the intraprotein and protein-solvent potential energy functions. An exchange of coordinates is allowed between neighboring replicas if the Monte Carlo metropolis criterion is satisfied.^41,42^ The HREMD method was chosen because it does not necessitate a predefined reaction coordinate. The advantage of HREMD over temperature replica exchange MD is that HREMD crosses entropic barriers^74^ more efficiently and a smaller number of replicas is sufficient, *i.e*. is computationally more efficient.

The total number of replicas (*n*) used, the scaling factor (*Λ_i_*) or the effective temperature (*T_i_*) of a replica and the average exchange probability (*p_eX_*) of the lowest rank replica are listed in **Tables S2-S4**. A *T_max_* of 400-450 K (lower limit) being needed, similar to *T_max_* = 400 K used in previous studies,^39,75,76^ and *p_ex_* ranging from 0.3 to 0.5. Moreover, to estimate the upper limit of effective temperature, we performed HREMD of Histatin 5 using a99SB-disp, *T_max_* = 800 K and 24 replicas (**Table S3**). This simulation generated the ensemble in the lowest rank replica similar to that of HREMD with *T_max_* = 450 K (**Fig. S11a**). However, we noted that replica from *T_i_* = 522 K and above sampled collapsed structures when compared to the ensemble of the lowest rank replica. Therefore, we suggest 450 *K*<*T_max_*<500 K is an appropriate choice for the upper limit of maximum effective temperature (**Fig. S11a**). However, choosing the higher value of *T_max_* would increase the number of replicas and thus computational cost.

In summary, we demonstrate HREMD simulations as an effective method to generate accurate structural ensembles of three IDPs with varying amino acid composition (Histatin 5, Sic 1 and SH4UD). The unbiased HREMD trajectories, calculated without using any experimental input or predefined reaction coordinate, are in excellent agreement with SAXS, SANS and NMR observables. Nonetheless, comparison to experimental data was imperative to confirm the accuracy of MD results. Moreover, HREMD simulations performed using two recent molecular mechanics force fields (a03ws and a99SB-disp) converge to the same distribution of *Rg*. In contrast, neither of the force fields could reproduce SAXS experiments with standard MD of the same cumulative length as HREMD. The results suggest adequately sampled simulations using recent IDP specific force fields can reliably generate the 3D ensembles of IDPs (**Fig. S12**), which is a prerequisite to an understanding of the biological function of IDPs. We also report that despite differences in their sequence, all three IDPs have similar local chain statistics for short lengths (less than ~ 10 residues). More studies are required to establish whether this is a universal IDP behavior.

## MATERIALS AND METHODS

### Experimental SAXS and NMR data

The experimental SAXS data of Histatin 5, Sic 1 and SH4UD were taken from Henriques et. al. (2015),^30^ Protein Ensemble Database (http://pedb.vib.be)^46^ and our previous work^40^ respectively. Similarly, NMR chemical shifts of backbone atoms, (*C^α^, C^β^, N^H^, H^α^, H^N^*) of Histatin 5, Sic 1 and SH4UD were acquired from the literature,^47^ Protein Ensemble Database^46^ and Biological Magnetic Resonance Data Bank (BMRB) database entry 15563^48^ respectively.

### MD simulations

The initial 3D structures of IDPs were obtained from I-TASSER.^77^ An MD-equilibrated starting structure with *R_g_* value close to experimental SAXS was chosen for the production simulation of each IDP. The same starting structure of IDP was utilized for each force field and sampling method.

We performed standard molecular dynamics simulations with two recently optimized force fields, Amber ff03ws^19,78,79^ with TIP4P/2005s^19^ (a03ws) and Amber ff99SB-*disp*^16,80^ with the modified TIP4P-D^16,21^ water model (a99SB-disp) using GROMACS.^81–86^ All bonds involving hydrogen atoms were constrained using LINCS algorithm.^87^ The Verlet leapfrog algorithm was used to numerically integrate the equation of motions with a time step of 2 fs. A cutoff of 1.2 nm was used for short-range electrostatic and Lennard-Jones interactions. Long-range electrostatic interactions were calculated by particle-mesh Ewald^88^ summation with a fourth order interpolation and a grid spacing of 0.16 nm. The solute and solvent were coupled separately to a temperature bath of 300 K, 293 K and 300 K for Histatin 5, Sic 1 and SH4UD respectively to match the temperatures measured at the experiments using modified Berendsen thermostat with a relaxation time of 0.1 ps. The pressure coupling was fixed at 1 bar using Parrinello-Rahman algorithm^89^ with a relaxation time of 2 ps and isothermal compressibility of 4.5*10^-5^ bar^-1^. The energy of each system was minimized using 1000 steepest decent steps followed by 1 ns equilibration at NVT and NPT ensembles. The production runs were carried out in the NPT ensemble.

### Enhanced sampling MD simulations

We employed replica-exchange with solute tempering 2 (REST2),^41,42^ a Hamiltonian Replica-Exchange MD (HREMD) simulation method to enhance the conformational sampling. REST2 is implemented in GROMACS^81–86^ patched with PLUMED.^90^ The interaction potentials of intraprotein and protein-solvent were scaled by a factor *Λ* and 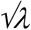 respectively, while water-water interactions were unaltered.^41,42,76,91^ The scaling factor *Λ*, and corresponding effective temperatures *T_i_* of the *i*^th^ replica are given by,

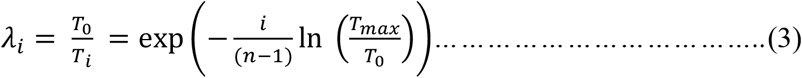

where *T_0_* and *T_max_* are the effective temperatures of lowest rank (unscaled) and the highest rank replicas respectively, and *n* is the total number of replicas used. For analysis we use only the trajectory of the unscaled for lowest rank replica (*λ_0_*=1 or *T_0_*). Exchange of coordinate between neighboring replicas was attempted every 400 MD steps. The details of HREMD and standard MD simulations are shown in **Tables S1-S4**. The secondary structure prediction was calculated with DSSP.^92^

### Error analysis

To estimate the error from HREMD trajectory, we divided the trajectory into five equal blocks each containing 10,000 frames (0-100, 100-200, 200-300, 300-400 and 400-500 ns). The mean value for each block, *m_i_* (*i*=1 to 5), was first calculated. The reported error bars are the standard error of the mean of the (*m_1_, m_2_, m_3_, m_4_, m_5_*) distribution.

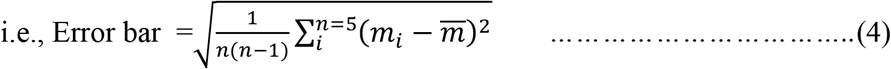

where 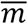 is the mean value and *n*=5 is the number of blocks used.

### Theoretical SAXS profiles

The theoretical SAXS and SANS intensities were calculated with SWAXS^45,93^ and SASSENA,^94^ respectively, by taking into account of explicit hydration water, which contributes to the signal.^45^ The agreement between experiment and simulation was determined by a *χ^2^* value:

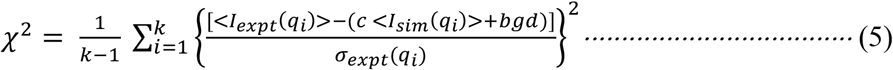

where <*I_expt_(q)*> and <*I_sim_(q)*> are the ensemble averaged experimental and theoretical SAXS data, respectively, *k* is the number of experimental *q* points, *c* is a scaling factor, *bgd* is a constant background and *σ_expt_* is the experimental error. In Eq. (4), *c* is a factor to scale calculated values to the experiment because the experimental values are often expressed in arbitrary units. It does not change the shape of the SAXS curve. Similarly, *bgd* is used to incorporate the uncertainty due to mismatch in buffer subtraction at higher *q*-values^13^ in the experiment.

### Theoretical NMR chemical shifts

Theoretical NMR chemical shifts were calculated with SHIFTX2^49^ by taking the average over all frames from the MD trajectory. The discrepancy between calculated and experimental values are measured by Mean Normalized Deviation is defined as,

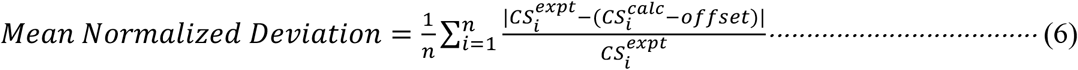

where 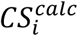 and 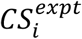 are the theoretical and experimental NMR chemical shift values respectively of residue index *i* of a protein with *n* number of residues, *offset* obtained from linear regression analysis are used for each backbone atom (*N^H^, C^α^, C^β^*) and IDP (Histatin 5, Sic 1, SH4UD) as shown in **Figs. S3-S5** and |…| is the modulus of the value enclosed.

## ACKNOWLEDGEMENTS

We thank Dr. Marie Skepö for generously sharing the SAXS data of Histatin 5. This research was supported by project ERKP300 funded by the Office of Biological & Environmental Research in the Department of Energy Office of Science. This research used a computing resource of the Oak Ridge Leadership Computing Facility (ALCC) and of the CADES at the Oak Ridge National Laboratory, which is supported by the Office of Science of DOE under Contract No. DEAC05-00OR22725.

