## Supplementary Information for "Full structural ensembles of intrinsically disordered proteins from unbiased molecular dynamics simulations"

### S1. Sequence and chemical properties of IDPs.

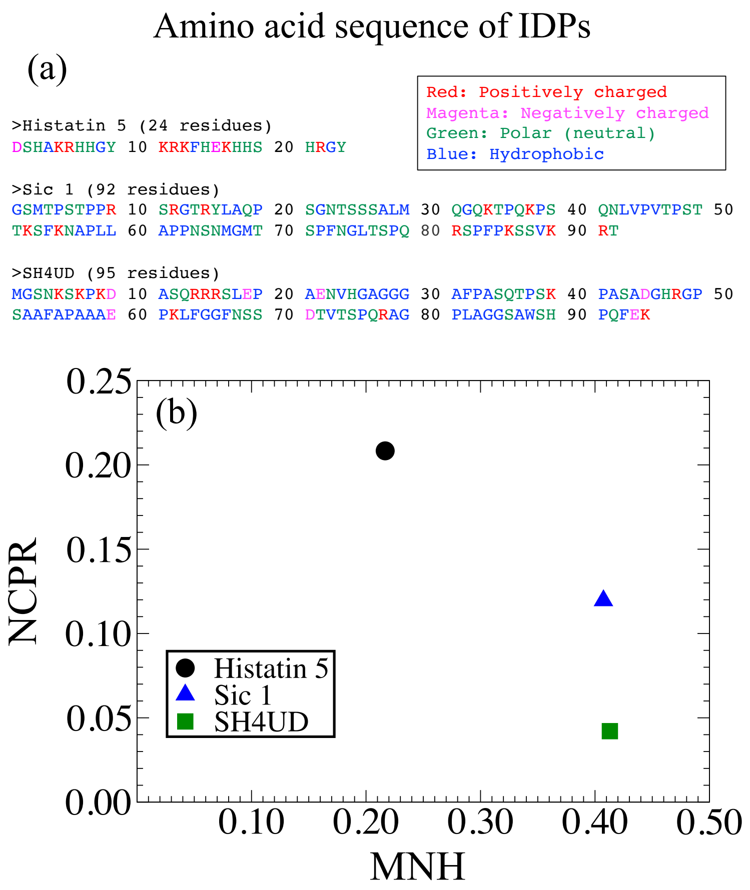

Fig. S1. (a) Sequence of the three IDPs (Histatin 5, Sic 1 and SH4UD) studied here. The red, magenta, green and blue colored letters represent positively charged, negatively charged, neutral polar and hydrophobic residues respectively. (b) Net charge per residue (NCPR) vs. mean normalized hydrophobicity (MNH) plot, also known as Uversky diagram is shown.

The sequence of Histatin 5, Sic 1 and SH4UD are shown in **Fig. S1a**. The net charge per residue (NCPR) of each protein sequence was calculated as follows,

$$NCPR = \frac{1}{n}\left|\sum_{i=1}^{n} q_i\right| \quad \text{(S1)}$$

where $q_i$ is the charge on $i^{th}$ residue and, $n$ is the total number of residues and |…| represents the absolute value. ExPASy[1] was used to calculate the normalized hydrophobicity of each protein residue by the Kyte and Doolittle approximation[2] with a window size of 5 residues and normalized to a scale of 0 to 1. The mean normalized hydrophobicity (MNH) is defined as,

$$MNH = \frac{1}{n}\sum_{i=1}^{n} H_i^{norm} \quad \text{(S2)}$$

where $H_i^{norm}$ is the normalized hydrophobicity of residue $i$ to a scale of 0 to 1 of each amino acid residue and $n$ is the total number of residues in a protein sequence. NCPR vs. MNH of each protein sequence is plotted in **Fig. S1b**. Sic 1 and SH4UD have nearly the same number of amino acid residues and MNH, but different NCPR.

### S2. MD simulation details.

The standard MD and HREMD simulations were conducted using two force fields, Amber ff03ws with TIP4P/2005s[3] water model (a03ws) and Amber ff99SB-*disp* with the TIP4P-D[4] water model (a99SB-disp).The details of the simulations are shown in the tables below.

Table S1. Details of standard MD simulations. Multiple copies were run with different initial velocity distributions.

| IDP | *T* (K) | Force field | Number of atoms | Number of copy x Length of simulation (ns) |
|---|---|---|---|---|
| Histatin 5 | 300 | a03ws | ~76k | 5x1000 |
| | | a99SB-disp | | |
| Sic 1 | 293 | a03ws | ~690k | 1x3000+3x1700 |
| | | a99SB-disp | | |
| SH4UD | 300 | a03ws | ~320k | 1x4000+5x1200 |
| | | a99SB-disp | | 1x4000+4x1600 |

Table S2. Details of HREMD simulations. $T_0$ is the effective temperature of the lowest-rank replica used in the analysis. $T_{max}$ is the effective temperature of the highest-rank replica. $p_{ex}$ is the average exchange probability of the lowest rank replica. Remarks indicate the agreement between the lowest rank replica ensemble with experimental data. The number of atoms in each simulation is the same as that in Table S1. The HREMD runs in bold-italics-large font are used to obtain the data shown in the main text. If a simulation did not agree with SAXS, we did not test its agreement with NMR.

| **IDP** | **Force field** | **$T_0$ (K)** | **$T_{max}$ (K)** | **Avg. $p_{ex}$ of $T_0$ replica** | **# of replicas (each 500 ns long)** |
|---|---|---|---|---|---|
| Histatin 5 | a03ws | 300 | ***425*** | ***0.4*** | ***10*** |
| | a99SB-disp | | ***450*** | ***0.3*** | ***10*** |
| | | | 800 | 0.3 | 24 |
| Sic 1 | a03ws | 293 | ***400*** | ***0.5*** | ***16*** |
| | a99SB-disp | | ***400*** | ***0.4*** | ***16*** |
| SH4UD | a03ws | 300 | ***400*** | ***0.6*** | ***20*** |
| | a99SB-disp | | ***450*** | ***0.5*** | ***20*** |

Table S3. Scaling factor $\lambda_i$- and temperature $T_i$- of the $i^{th}$ replica for HREMD simulations of IDPs.

| **Histatin 5** | | | | | |
|---|---|---|---|---|---|
| **$T_0$=300 K - $T_{max}$=425 K** | | **$T_0$=300 K - $T_{max}$=450 K** | | **$T_0$=300 K - $T_{max}$=800 K** | |
| $\lambda_i$ | $T_i$ (K) | $\lambda_i$ | $T_i$ (K) | $\lambda_i$ | $T_i$ (K) |
| 1.00 | 300.00 | 1.00 | 300.00 | 1.00 | 300.00 |
| 0.96 | 311.84 | 0.96 | 313.82 | 0.96 | 313.07 |
| 0.93 | 324.14 | 0.91 | 328.29 | 0.92 | 326.71 |
| 0.89 | 336.93 | 0.87 | 343.41 | 0.88 | 340.94 |
| 0.86 | 350.23 | 0.84 | 359.24 | 0.84 | 355.80 |
| 0.82 | 364.05 | 0.80 | 375.79 | 0.81 | 371.30 |
| 0.79 | 378.41 | 0.76 | 393.11 | 0.77 | 387.47 |
| 0.76 | 393.35 | 0.73 | 411.23 | 0.74 | 404.36 |
| 0.73 | 408.87 | 0.70 | 430.18 | 0.71 | 421.97 |
| 0.71 | 425.00 | 0.67 | 450.00 | 0.68 | 440.36 |
| | | | | 0.65 | 459.54 |
| | | | | 0.63 | 479.56 |
| | | | | 0.60 | 500.46 |
| | | | | **0.57** | **522.26 ($T_{collapse}$)** |
| | | | | 0.55 | 545.01 |
| | | | | 0.53 | 568.76 |
| | | | | 0.51 | 593.54 |
| | | | | 0.48 | 619.40 |
| | | | | 0.46 | 646.38 |
| | | | | 0.44 | 674.54 |
| | | | | 0.43 | 703.93 |
| | | | | 0.41 | 734.60 |
| | | | | 0.39 | 766.60 |
| | | | | 0.38 | 800.00 |

| **Sic 1** | | **SH4UD** | | | |
|---|---|---|---|---|---|
| **$T_0$=293 K - $T_{max}$=400 K** | | **$T_0$=300 K - $T_{max}$=400 K** | | **$T_0$=300 K - $T_{max}$=450 K** | |
| $\lambda_i$ | $T_i$ (K) | $\lambda_i$ | $T_i$ (K) | $\lambda_i$ | $T_i$ (K) |
| 1.00 | 293.00 | 1.00 | 300.00 | 1.00 | 300.00 |
| 0.98 | 299.14 | 0.98 | 304.58 | 0.98 | 306.47 |
| 0.96 | 305.42 | 0.97 | 309.22 | 0.96 | 313.08 |
| 0.94 | 311.82 | 0.96 | 313.94 | 0.94 | 319.83 |
| 0.92 | 318.36 | 0.94 | 318.73 | 0.92 | 326.73 |
| 0.90 | 325.04 | 0.93 | 323.59 | 0.90 | 333.78 |
| 0.88 | 331.85 | 0.91 | 328.53 | 0.88 | 340.98 |
| 0.86 | 338.81 | 0.90 | 333.54 | 0.86 | 348.33 |
| 0.85 | 345.92 | 0.89 | 338.63 | 0.84 | 355.85 |
| 0.83 | 353.17 | 0.87 | 343.80 | 0.83 | 363.52 |
| 0.81 | 360.58 | 0.86 | 349.04 | 0.81 | 371.36 |
| 0.80 | 368.14 | 0.85 | 354.37 | 0.79 | 379.38 |
| 0.78 | 375.86 | 0.83 | 359.77 | 0.77 | 387.56 |
| 0.76 | 383.74 | 0.82 | 365.26 | 0.76 | 395.92 |
| 0.75 | 391.78 | 0.81 | 370.84 | 0.74 | 404.46 |
| 0.73 | 400.00 | 0.80 | 376.49 | 0.73 | 413.18 |
| | | 0.78 | 382.24 | 0.71 | 422.09 |
| | | 0.77 | 388.07 | 0.70 | 431.20 |
| | | 0.76 | 393.99 | 0.68 | 440.50 |
| | | 0.75 | 400.00 | 0.67 | 450.00 |

Table S4. The ensemble-average values of $R_g$ from standard MD and HREMD simulations.

| **IDP** | **FF** | ***$T_0$* (K)** | **$R_{g,MD}$ (nm)** | **HREMD *$T_{max}$* (K)** | **$R_{g,HREMD}$ (nm)** |
|---|---|---|---|---|---|
| Histatin 5 | a03ws | 300 | 1.10±0.03 | 425 | 1.23±0.02 |
| | a99SB-disp | | 1.15±0.03 | 450 | 1.26±0.01 |
| | | | | 800 | 1.24±0.01 |
| Sic 1 | a03ws | 293 | 3.04±0.04 | 400 | 3.14±0.13 |
| | a99SB-disp | | 2.60±0.09 | 400 | 3.05±0.04 |
| SH4UD | a03ws | 300 | 2.05±0.14 | 400 | 2.60±0.05 |
| | a99SB-disp | | 2.00±0.07 | 450 | 2.50±0.10 |

### S3. Small-angle neutron scattering (SANS) of SH4UD.

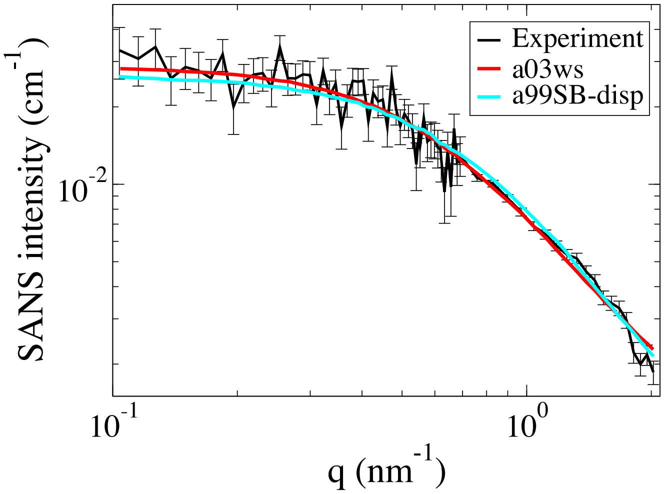

Fig. S2. Experimental (black) and simulation-derived small-angle neutron scattering (SANS) profiles of SH4UD are compared. The theoretical SANS data are calculated taking into account of explicit hydration shell around IDP using software SASSENA[5] from HREMD simulations of a03ws (red) and a99SB-disp (cyan) force fields.

### S4. Quality of agreement between small-angle scattering experiments and MD simulations.

The quality of agreement of theoretical SAXS and SANS profiles to experimental data is quantified by calculating chi-square ($\chi^2$) using Eq. (5). The values of $\chi^2$ for IDP are listed in Table S5.

Table S5. $\chi^2$, defined in **Eq. (5)**, between small-angle scattering experiments and HREMD simulations averaged over the entire simulation trajectory.

| | | $\chi^2$ | | |
|---|---|---|---|---|
| Sampling method | Force field | Histatin 5 (SAXS) | Sic 1 (SAXS) | SH4UD (SAXS, SANS) |
| Standard MD | a03ws | 5.3 | 0.2 | 6.2 |
| | a99SB-disp | 3.0 | 3.4 | 6.8 |
| HREMD | a03ws | 2.0 | 0.2 | 1.2, 1.3 |
| | a99SB-disp | 2.0 | 0.2 | 1.0, 1.3 |

### S5. Comparison of experimental and calculated NMR chemical shifts of backbone atoms in IDPs.

The linear regression analysis is shown to compare the agreement between experimental and MD-calculated chemical shifts of IDPs. The *offset* values obtained from linear fit are used in Eq. (6) to calculate mean normalized deviation.

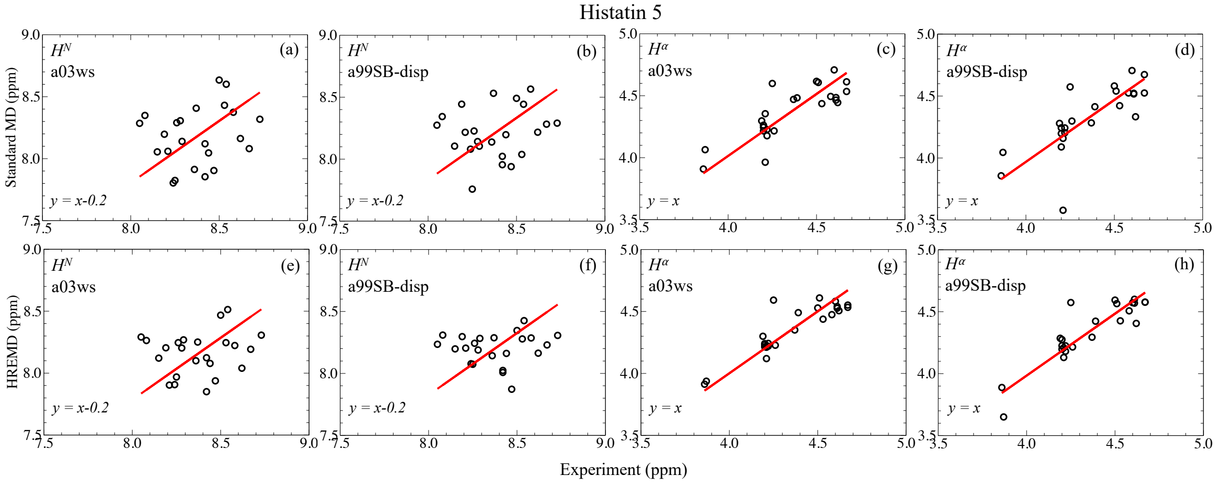

Fig. S3. Comparison between the ensemble-averaged calculated and experimental NMR chemical shifts of backbone atoms ($H^N$ and $H^\alpha$) of Histatin 5.

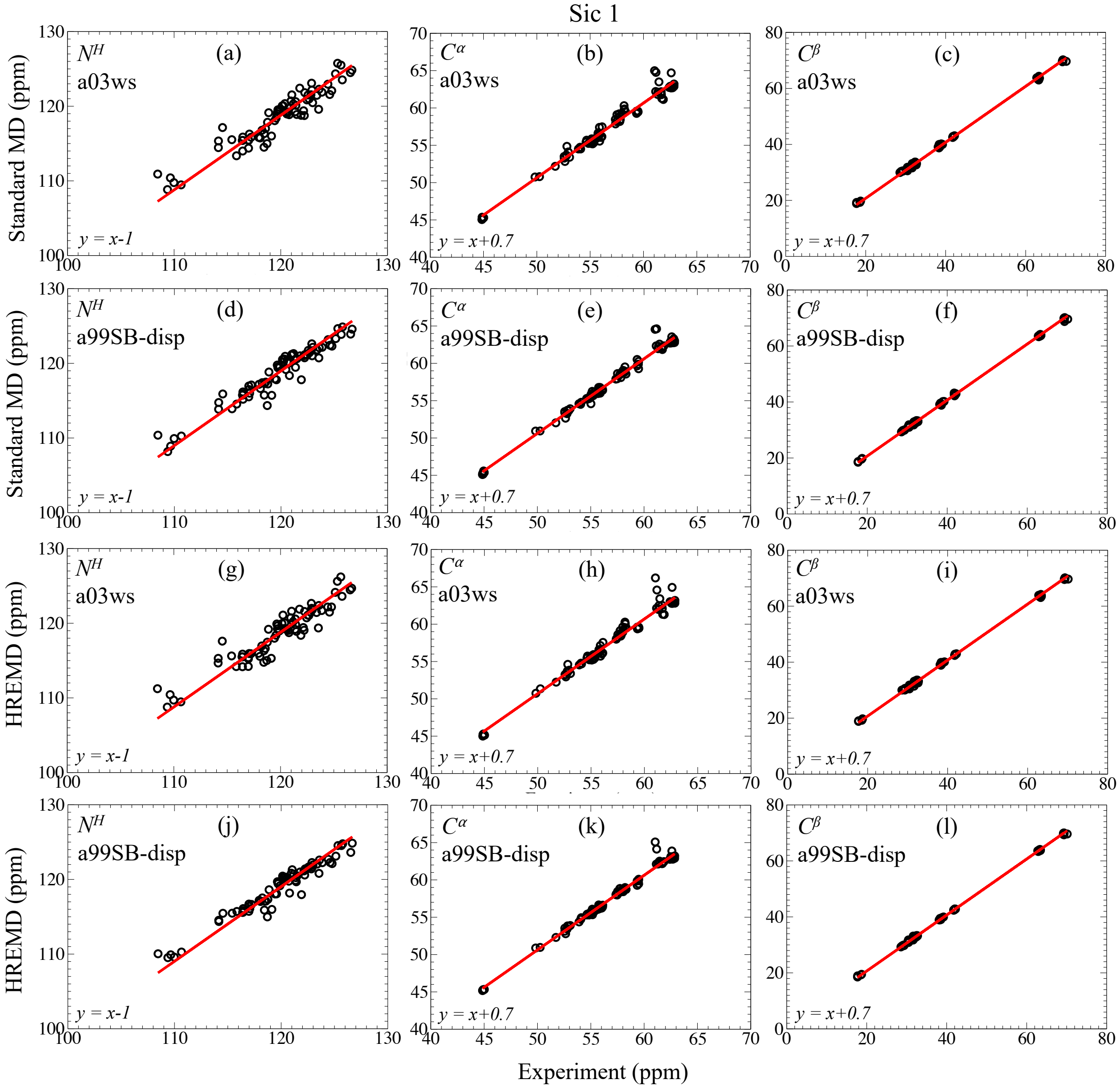

Fig. S4. Comparison between the ensemble-averaged calculated and experimental NMR chemical shifts of backbone atoms ($N^H$, $C^\alpha$, $C^\beta$) of Sic 1.

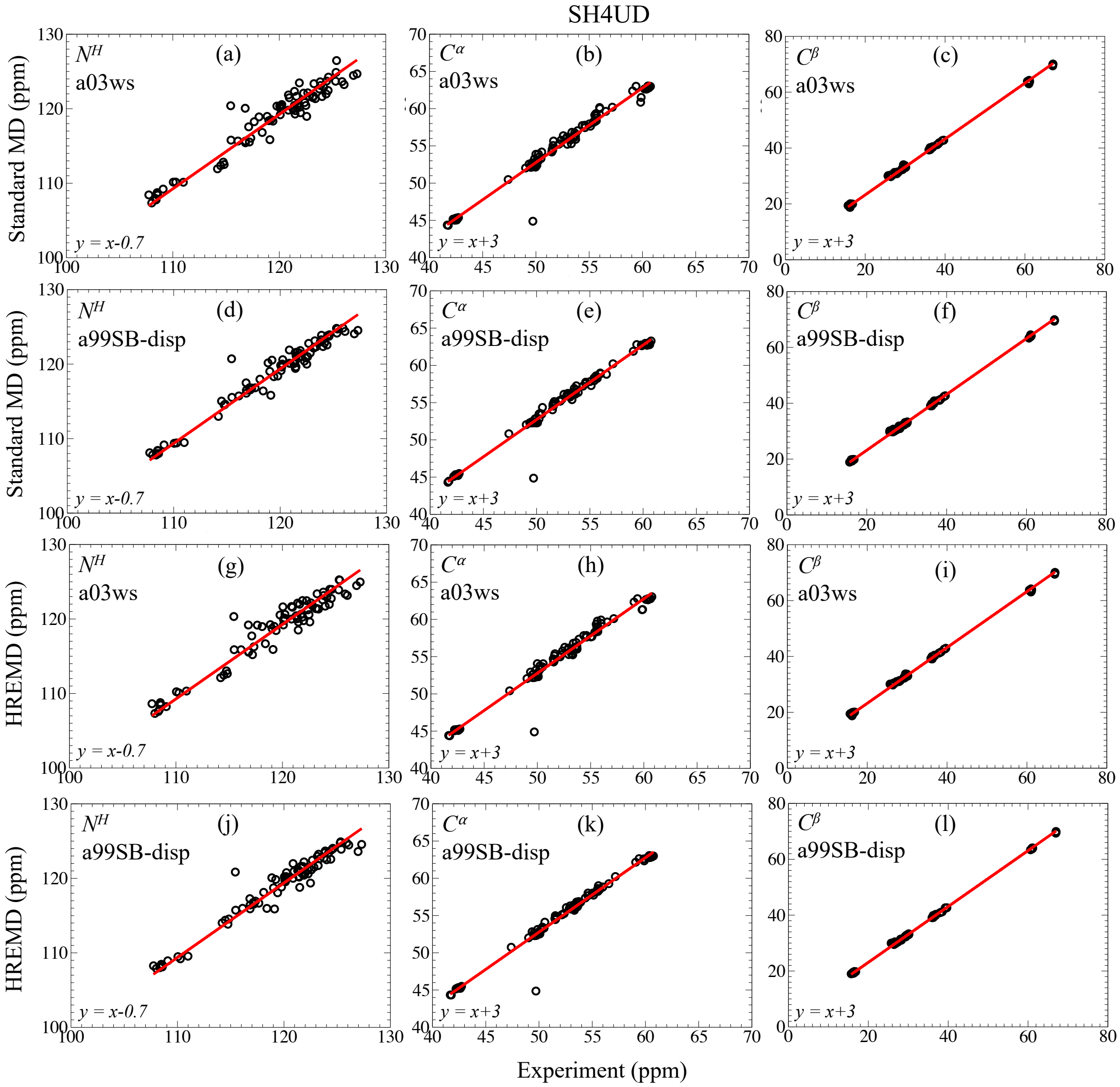

Fig. S5. Comparison between the ensemble-averaged calculated and experimental NMR chemical shifts of backbone atoms ($N^H$, $C^\alpha$, $C^\beta$) of SH4UD.

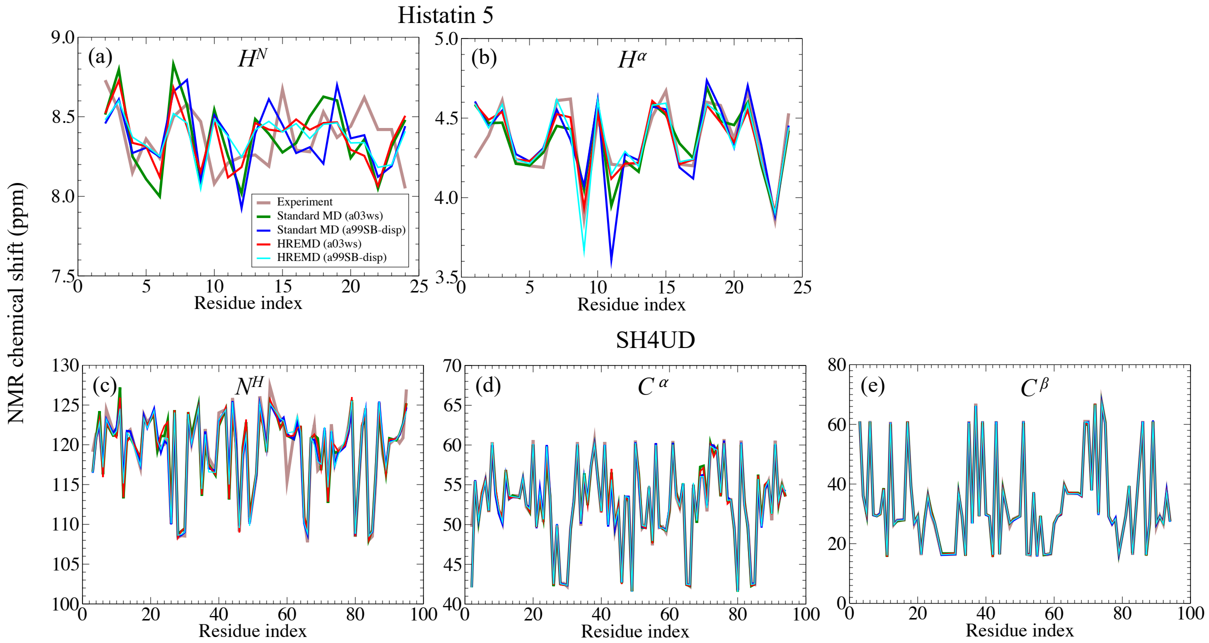

Fig. S6. Comparison between the ensemble-averaged calculated and experimental NMR chemical shifts of the Histatin 5 backbone atoms (a) $H^N$ and (b) $H^\alpha$ and SH4UD backbone atoms (c) $N^H$, (d) $C^\alpha$ and (e) $C^\beta$; and. The experimental[6,7] and calculated values from MD simulations are shown by different color lines.

### S6. Propensity of coil and secondary structures in IDPs.

Histatin 5, Sic 1 and SH4UD are shown to possess transient $3_{10}$- and α- helices, but negligible β-sheets (**Fig. S4 and S5**). The lack of β-sheet structure may explain because it requires the formation of hydrogen bond between residues distant in sequence space, which may be unlikely to form in IDPs which adopt extended conformation and also confirmed by the lack of long-range contacts (**Fig. S9**).

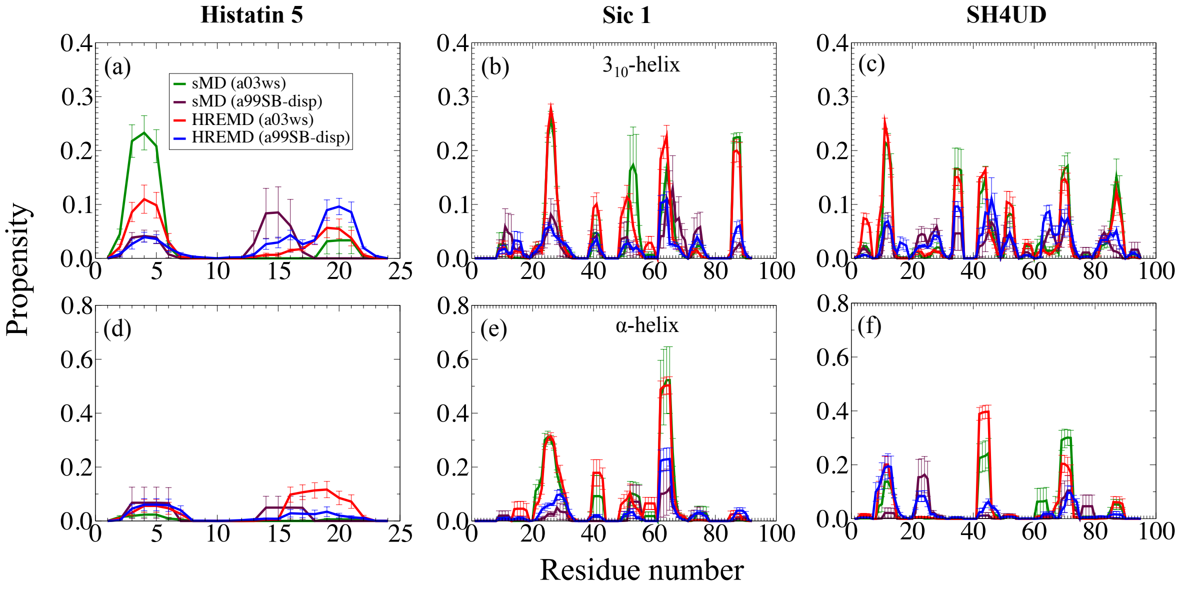

Fig. S7. Helical structure propensity. Propensity is scaled between 0 and 1 implying none (0%) and all the snapshot (100%) recorded in MD trajectory respectively here and in the subsequent figures below.

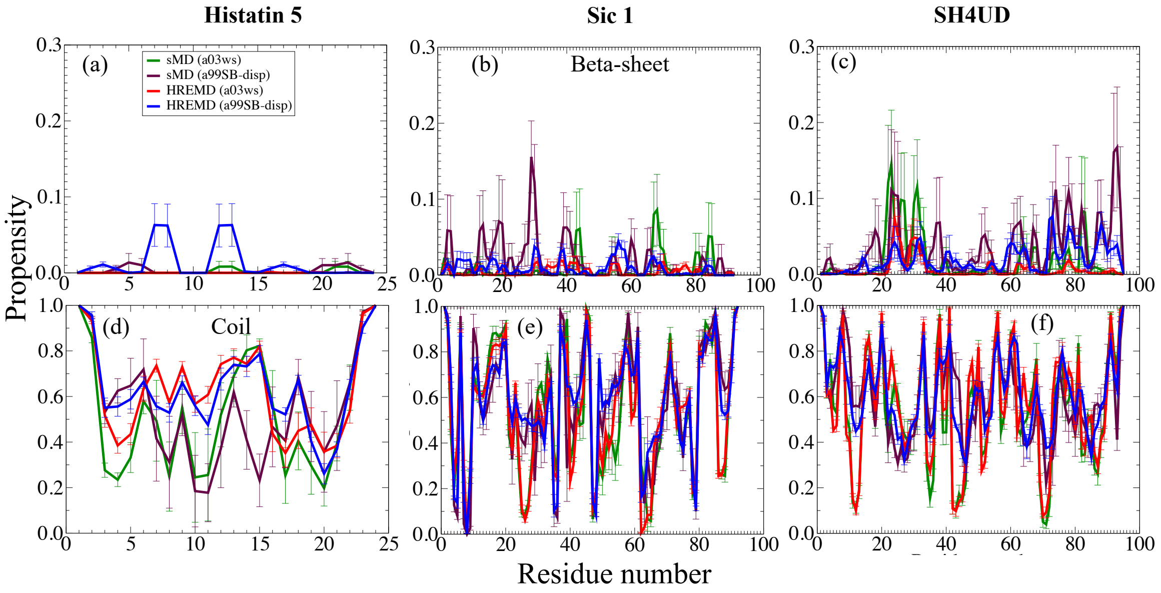

Fig. S8. Propensity of β-sheet and coil structures.

### S7. Contact maps of IDPs.

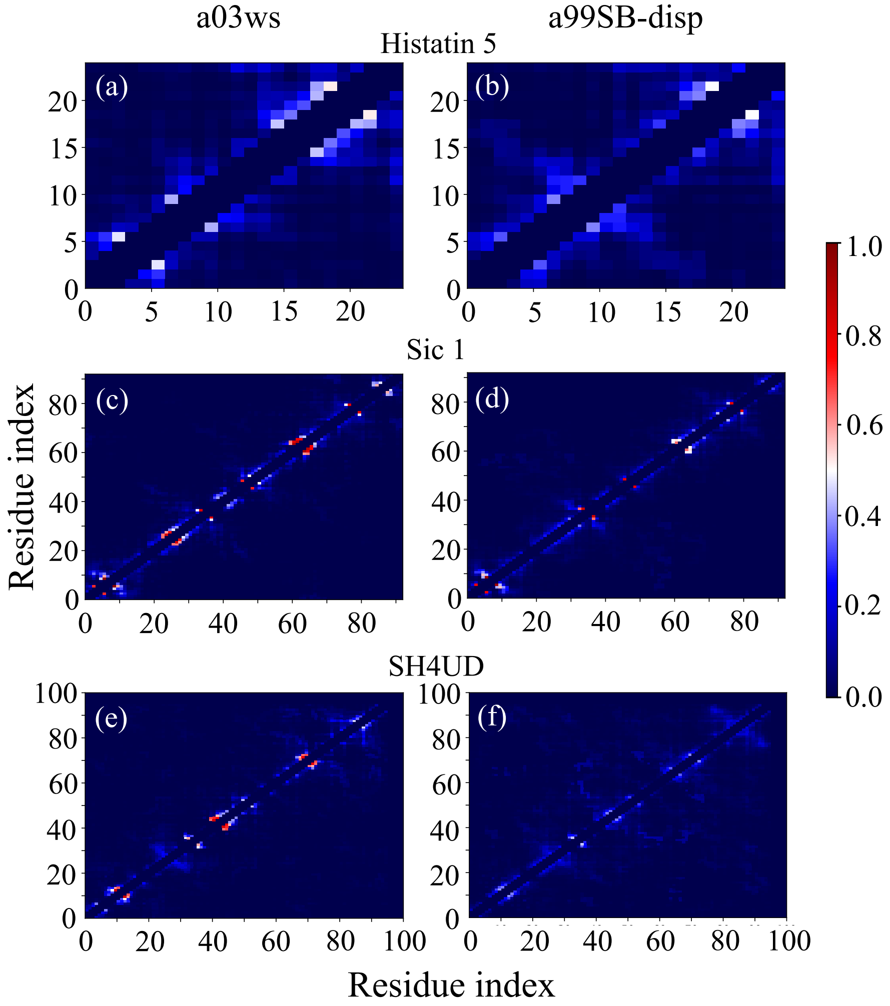

Fig. S9. Contact maps of IDPs calculated using Contact Map Explorer (https://contact-map.readthedocs.io/en/latest/index.html). The cut-off distance of 0.45 nm was used and the atoms of 2 residues on either side of given residue and to itself are excluded. The color index represents the fraction of native contacts in HREMD trajectory for each simulation.

### S8. Convergence of HREMD trajectories with respect to SAXS experiments.

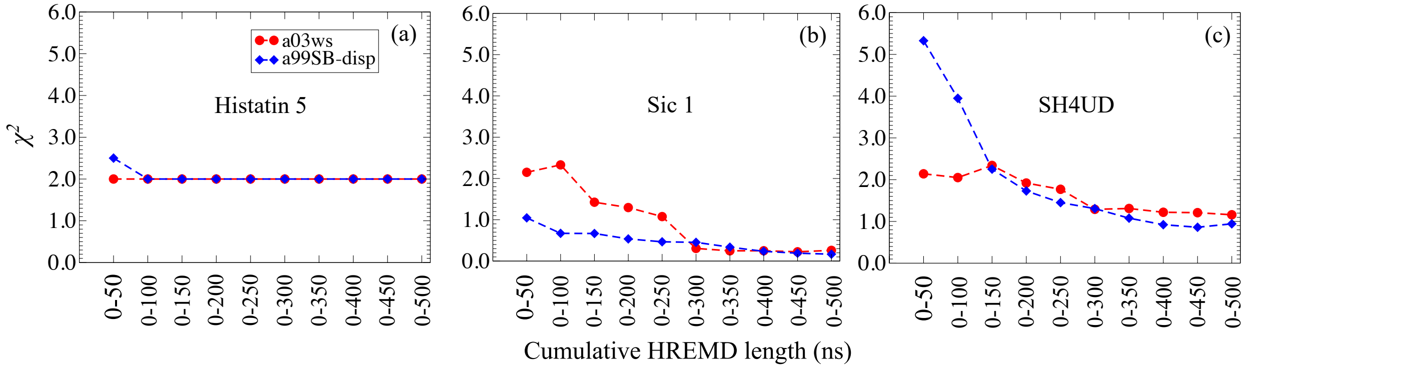

Fig. S10. $\chi^2$ values quantifying the agreement between theoretical vs. experimental SAXS data (**Eq. 5**) with respect to cumulative length of HREMD simulations.

### S9. HREMD of Histatin 5 with $T_{max}$ = 800 K.

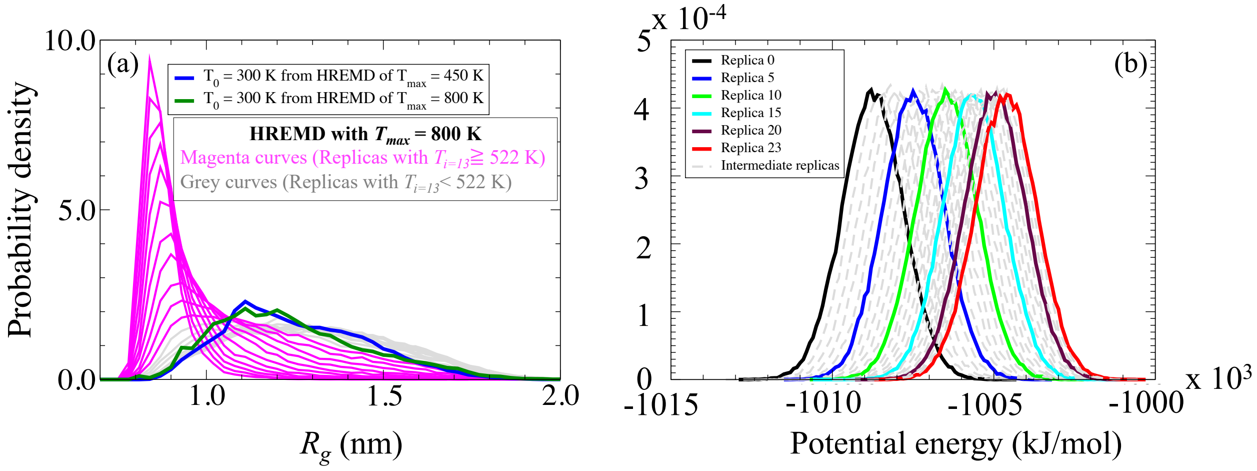

Fig. S11. Results from all the replicas of HREMD simulations of Histatin 5 (a99SB-disp) using $T_{max}$ = 800 K. (a) Histograms of $R_g$ of with temperatures above $T_{i=13}$ = 522 K (magenta) generate overly collapsed conformations compared below $T_{i=13}$ (grey). The lowest rank replica ($T_0$ = 300 K) from HREMD using $T_{max}$ = 800 K is shown in green. For comparison, we show the lowest ranked replica from the $T_{max}$ = 450 K HREMD reported in the main text with blue curve. (b) Potential energy of a system for all the replicas from HREMD with $T_{max}$ = 800 K. A overlap between neighboring replicas confirms that there is no phase transition.

### S10. Ensemble of 3D structures of protein from HREMD simulations.

(a) **Histatin 5**

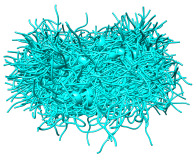

(b) **Sic 1**

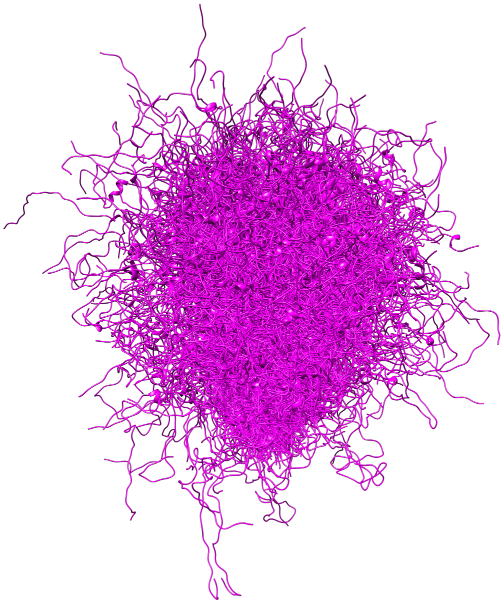

(c) **SH4UD**

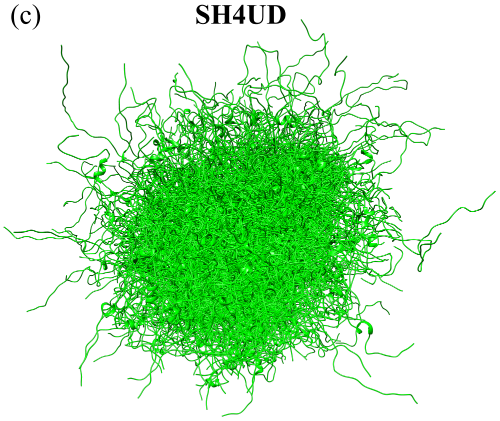

Fig. S12. Ensemble of structures of proteins from HREMD simulations. Ensemble obtained from structures saved every 500 ps (total frames 1k) of lowest rank replica of 500 ns long HREMD trajectory (a99SB-disp) of (a) Histatin

5, (b) Sic 1, and (c) SH4UD. The size of the ensemble is not drawn to scale and thus cannot be compared to each other.
